# Myosin II Tension Sensors Visualize Force Generation within the Actin Cytoskeleton in Living Cells

**DOI:** 10.1101/623249

**Authors:** Ryan G. Hart, Divya Kota, Fangjia Li, Diego Ramallo, Andrew J. Price, Karla L. Otterpohl, Steve J. Smith, Alexander R. Dunn, Jing Liu, Indra Chandrasekar

## Abstract

Type II myosin motors generate cytoskeletal forces that are central to cell division, embryogenesis, muscle contraction, and many other cellular functions. However, at present there is no method that can directly measure the forces generated by myosins in living cells. Here we describe a Förster resonance energy transfer (FRET)-based tension sensor that can measure forces generated by Nonmuscle Myosin IIB (NMIIB) in living cells with piconewton (pN) sensitivity. Fluorescence lifetime imaging microscopy (FLIM)-FRET measurements indicate that the forces generated by NMIIB exhibit significant spatial and temporal heterogeneity, with inferred tensions that vary widely in different regions of the cell. This initial report highlights the potential utility of myosin-based tension sensors in elucidating the roles of cytoskeletal contractility in a wide variety of contexts.

## Introduction

Force generation by myosin II isoforms is fundamental to the function of all eukaryotic cells. In addition to the well-known role of myosin II in generating muscle contraction, nonmuscle myosin II (NMII) isoforms are responsible for generating the mechanical forces that power cell division, cell motility, membrane remodeling during endo/exocytosis and embryo morphogenesis (Betapudi, 2014; Chandrasekar et al., 2014; Milberg et al., 2017; Sellers, 2000; Vicente-Manzanares et al., 2009). Dysregulation of force generation by nonmuscle myosin II can result in or contribute to disease conditions that include muscular dystrophy, cardiomyopathy, and metastatic cancer (Sellers, 2000). In humans, cells express at least one of three known nonmuscle myosin II isoforms (IIA, IIB, IIC), which predominantly assemble into bipolar filament assemblies containing approximately 15-30 molecules of NMII (Billington et al., 2013). These bipolar filaments use energy generated by ATP hydrolysis to translocate actin filaments, and in so doing generate contractile forces. Recent advances demonstrate that both the localization and activation of NMII is subject to complex regulation (Beach et al., 2017; Beach and Hammer, 2015; Beach et al., 2014; Fenix et al., 2016; Hu et al., 2017). However, despite its biological importance, a method for directly visualizing the mechanical forces generated by NMII in living cells is presently lacking.

In vitro measurements have shown that single myosin catalytic domains generate forces of 4-5 pN (Finer et al., 1994). Recently developed Förster resonance energy transfer (FRET)-based tension sensors are well suited to investigate how motor proteins generate and propagate pN-scale forces in cells (Freikamp et al., 2016; Grashoff et al., 2010; Guo et al., 2014; Meng et al., 2008). One such sensor, termed the tension sensor module (TSMod), consists of an extensible flagelliform linker peptide flanked by FRET donor and acceptor fluorophores (Grashoff et al., 2010). Tension on TSMod results in the extension of the flagelliform linker and a corresponding decrease in FRET efficiency that can be quantified to yield an estimate of the magnitude of the force (i.e. tension) experienced by the module (Fig. 1A). FRET-based tension sensors have enabled the measurement of traction forces at integrin-based cell-matrix adhesions (Kumar et al., 2016; LaCroix et al., 2018; Nordenfelt et al., 2016), the nuclear LINC complex (Arsenovic et al., 2016), cell-cell adhesions (Borghi et al., 2012; Lagendijk et al., 2017; Price et al., 2018), and within the spectrin cytoskeleton (Krieg et al., 2014). FRET-based tension sensors likewise allow the measurement of mechanical tension on the host protein in intact, living model organisms (Krieg et al., 2014; Lagendijk et al., 2017; Meng et al., 2008; Yamashita et al., 2016).

**Figure 1:**
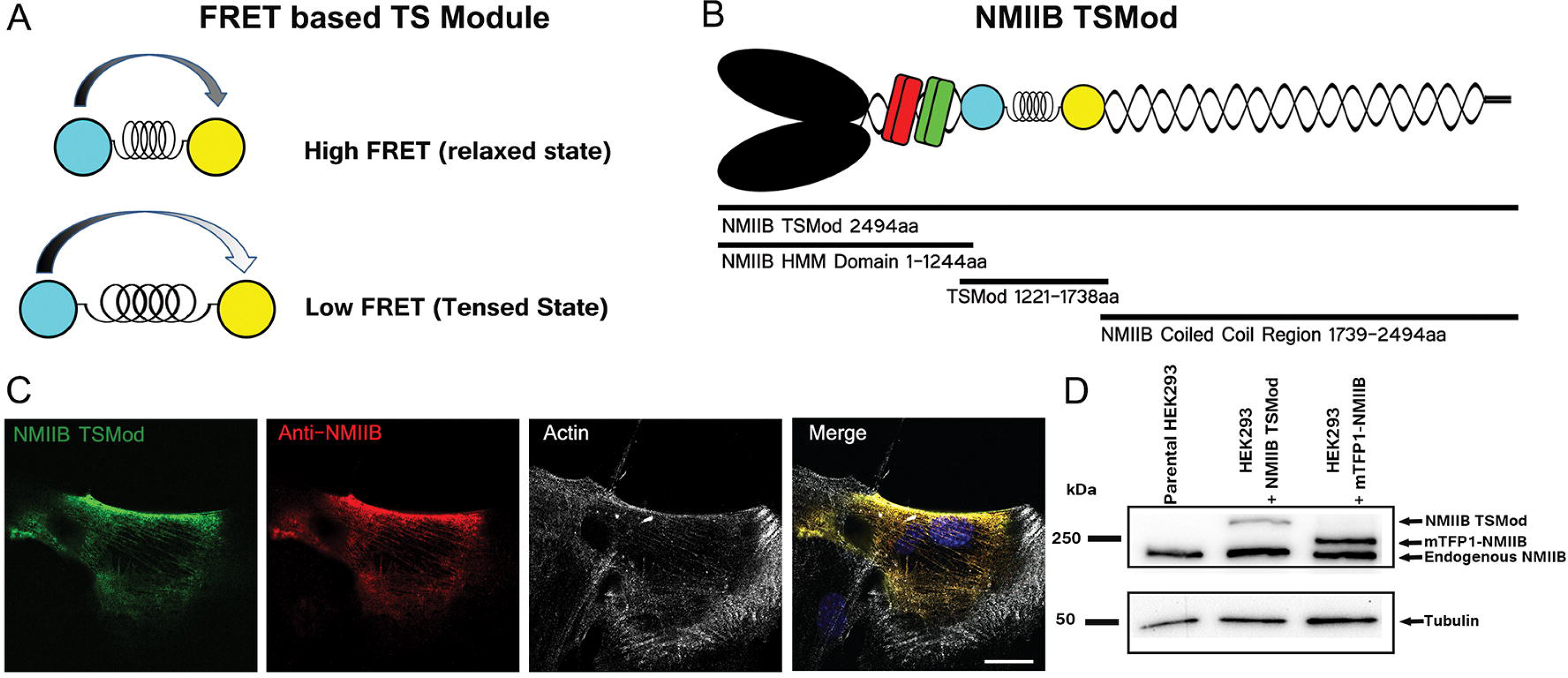
Design and development of a NMII tension sensor. Schematic of the FRET-based tension sensor module (TSMod) in its relaxed and tensed states (A). Cartoon illustrating the structural domains of NMIIB TSMod (B). NMIIB−/− mouse embryonic fibroblasts transiently expressing NMIIB TSMod (green) show NMIIB TSMod filaments are stained with an anti-NMIB antibody (red) and colocalize with actin filaments (white) (C). Immunoblot of parental HEK293 cells and cells expressing NMIIB TSMod or mTFP1-Donor only control, showing endogenous and exogenous protein bands (D), Scale: 10 μm.

FRET-based methods have also been widely applied to study conformational dynamics in motor proteins, including Myosin II (Iwai and Uyeda, 2008; Kasprzak, 2007; Muretta et al., 2015; Shih et al., 2000). Most of these studies were performed *in vitro* using purified, partial domains of skeletal (Muretta et al., 2015), smooth muscle or *Dictyostelium* (Shih et al., 2000; Zeng et al., 2006) myosin II (eg: heavy meromyosin or S1 fragment). *In vitro* optical tweezer microscopy experiments have also revealed details of the mechanism by which myosin II generates force (Finer et al., 1994; Spudich, 2001). Conversely, traction force microscopy and laser microdissection approaches can provide estimates for actomyosin-generated forces in intact cells (Polacheck and Chen, 2016; Sugita et al., 2011). However, to our knowledge a direct measurement of NMII-associated forces within the cytoskeleton has not been reported.

In this study, we developed NMIIB tension sensors that report on myosin-generated tension within living cells, with submicron and sub-second spatial and temporal resolutions. Our measurements reveal that actomyosin-associated forces are dynamic and exhibit spatial and temporal fluctuations in different subcellular regions in living cells. We anticipate that NMII-based tension sensors may be widely useful for measuring myosin-generated forces in living cells, and in future studies in intact organisms.

## Results

### Design and Generation of Nonmuscle Myosin IIB (NMIIB) Tension Sensors

NMII is a hexameric motor protein comprised of two heavy chains, each of which bind a regulatory and essential light chain (Fig.1B). The heavy chain is composed of the globular head, which contains both the actin-and ATP-binding sites and a coiled-coil tail domain, which mediates both dimerization and filament assembly (Sellers and Knight, 2007). To generate the NMIIB tension sensor, we inserted TSMod (Fig.1A) between amino acids A1220 and L1222, twenty amino acids N-terminal of the protease recognition site whose cleavage generates heavy meromyosin (a functional motor domain of NMII) (Sellers et al., 1981). We predicted that this region of myosin II might be conducive to manipulation without affecting the major structural regions of the heavy chain involved in ATPase activity, light chain binding, and bipolar filament assembly.

To evaluate the expression level and localization profile of NMIIB TSMod, we performed transient transfections of NMIIB −/− (null) mouse embryonic fibroblasts (MEFs) and stained for NMIIB and actin filaments (Fig.1C). We observed NMIIB TSMod organized into filaments along the cell body and trailing edge that co-localized with the actin cytoskeleton in migrating MEFs (Fig.1C). In human embryonic kidney (HEK293) cells, which express endogenous NMIIB, we observed the integration of NMIIB TSMod with the endogenous NMIIB in (Fig. S1G). Immunoblot analysis performed using HEK293 cells transfected with NMIIB TSMod or an N-terminally tagged, donor-only construct (Fig S1A; mTFP1-NMIIB) showed expression of exogenous mTFP1-NMIIB (255 kDa) or NMIIB TSMod (285 kDa) at levels comparable to endogenous NMIIB (228k Da) (Fig.1D). In addition, we generated control constructs that lacked a functional FRET donor or acceptor fluorophore, termed NMIIB TSMod T (mTFP1 G73S) and the NMIIB TSMod V (Venus G67S), respectively (Fig. S1B, C). Finally, we generated a construct in which the elastic linker peptide in the TSMod was replaced with a five amino acid (SGKRS) linker that holds the fluorophores in close proximity, yielding a construct whose FRET efficiency is minimally sensitive to tension (Fig. S1D). To probe the dependence of the TSMod insertion site on the force readout, we designed an alternative NMIIB tension sensor in which TSMod is inserted after the S1 fragment (1-844aa), and is hence named NMIIB-S1 TSMod (Fig. S1E).

### Characterization of NMIIB tension sensors and control constructs

We used Human Embryonic Kidney cells (HEK293) and osteosarcoma cells (U2OS) as initial model systems to evaluate the ability of NMIIB tension sensors to measure actomyosin-associated forces. FRET values were measured using fluorescence lifetime imaging microscopy (FLIM) with time-correlated single photon counting (TCSPC) in living cells. This method provides the gold-standard FRET efficiency measurements. Images were processed using PicoQuant software (Symphotime) to determine mTFP1 excited state lifetimes and hence FRET efficiency values along actomyosin filaments, which we use here as a general term for structures that appear to be stress fibers (Fig. S2). In addition to FLIM-FRET images of the whole cell, we also performed point measurements by selecting 10-30 ~0.3×0.3 μm regions of interest (ROIs) on actomyosin filaments to obtain localized, time-resolved FLIM-FRET time series with pixel dwell times of 0.2-0.3 ms, which is not possible when scanning the whole cell (Fig. S2A). Unless specified otherwise, point measurement ROIs were randomly selected along actomyosin filament after scanning a candidate cell.

As anticipated, in HEK293 cells the donor-only control mTFP1-NMIIB showed an average lifetime of 2.7 ± 0.1 ns (mean ± S.D.) corresponding to a FRET efficiency of 3 ± 4% (mean ± S.D.) i.e. indistinguishable from zero (Fig. 2A, A’, E & F). The NMIIB tension sensor, NMIIB TSMod, yielded an average mTFP1 lifetime of 2.3 ± 0.2 ns (mean ± S.D.) and a corresponding FRET efficiency of 15 ± 8% (mean ± S.D.) (Fig. 2B, B’, E & F). This FRET value is consistent with the presence of mechanical load, as it is appreciably less than the FRET efficiency expected in the absence of load (Grashoff et al., 2010; Rothenberg et al., 2018). The average NMIIB-5AA mTFP1 lifetime was 1.8 ± 0.2 ns with a corresponding FRET efficiency of 34 ± 9% (mean ± S.D.) (Fig. 2C, C’, E & F). NMIIB TSMod V, which lacks a functional FRET acceptor, yielded an average mTFP1 lifetime of 2.6 ± 0.1 ns, similar to that of mTFP1-NMIIB, and a corresponding FRET efficiency of 4 ± 5% (mean ± S.D.) (Fig. 2D, D’, E & F).

**Figure 2:**
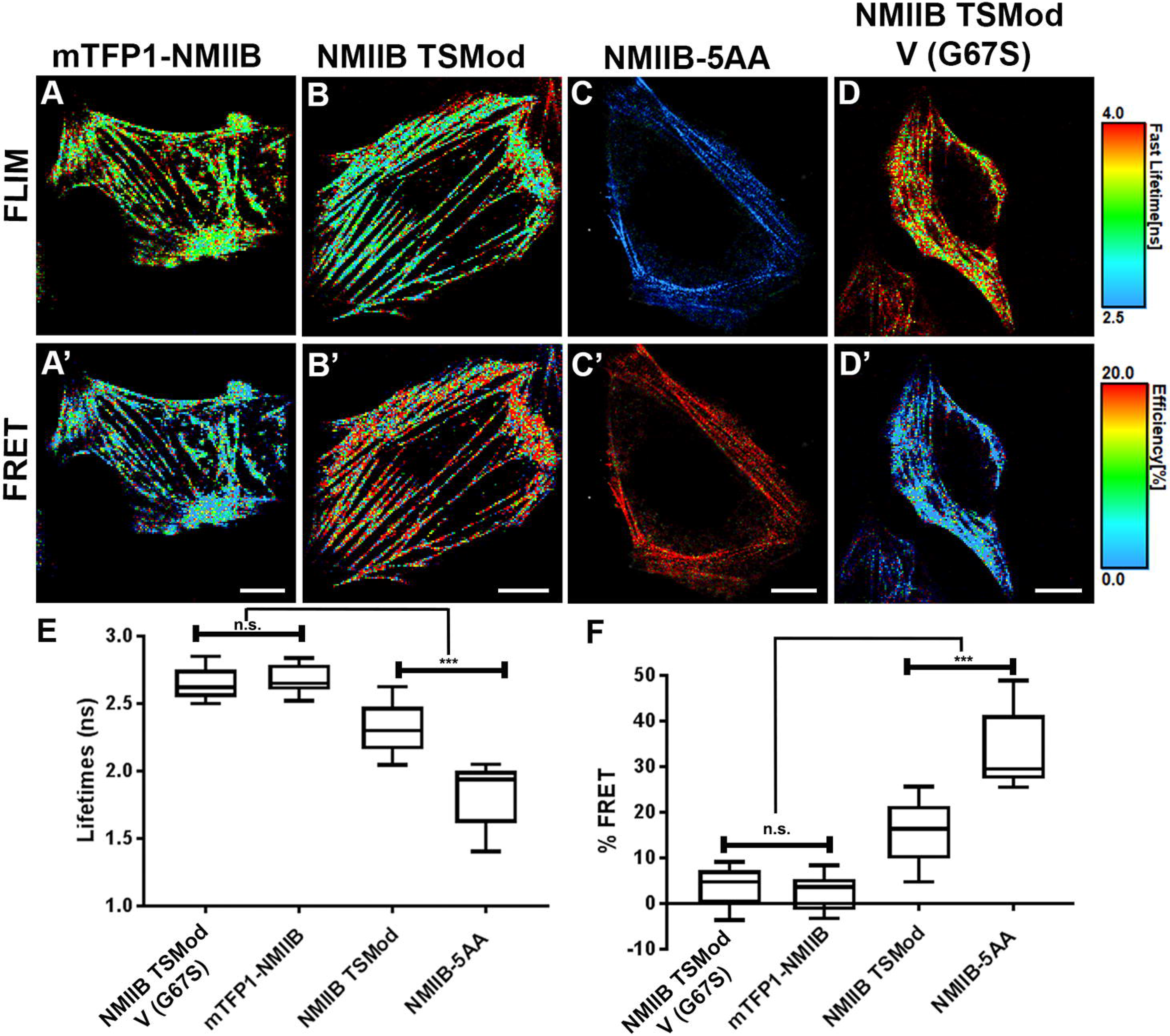
NMIIB TSMod FLIM-FRET measurements within the actomyosin filaments in HEK293 cells. Representative images from time correlated FLIM (A-D) and the corresponding FRET percentages (A’-D’) of HEK293 cells expressing NMIIB TSMod and control constructs. Scale: FLIM (2.5 – 4 ns) and FRET (0 – 20%). ROI measurements were performed along the actomyosin filaments in living cells expressing mTFP1-NMIIB [donor only control (A, A’, E and F)] and the NMIIB TSMod V (G67S) [dark acceptor control (D, D’, E and F)] show higher lifetimes (ns) and lower FRET (%) compared to NMIIB TSMod (B, B’, E and F). NMIIB-5AA shows low lifetimes and high FRET efficiency (C, C’, E and F). n=15-20 cells. ROI FLIM-FRET were measurements extracted from point measurements and whole cell images using PicoQuant SymphoTime software, and were pooled from 3-5 separate experimental sets. Statistical analyses using Kruskal-Wallis test with multiple comparisons show significant differences (P value < 0.0001) between all parameters, except between controls NMIIB TSMod G67S and mTFP1-NMIIB, Scale: 10 μm.

We also expressed NMIIB TSMod in U2OS cells and recorded an average lifetime of 2.3 ± 0.2 ns, corresponding to an average FRET efficiency of 18 ± 9% (mean ± S.D.) (Fig. S3). In these cells the FRET efficiencies of the control constructs, NMIIB TSMod V (G67S), mTFP1-NMIIB, and NMIIB-5AA were 4 ± 5%, 5 ± 5%, and 42 ± 3% (mean ± S.D.) respectively (Fig. S3). Thus, FRET measurements were consistent across the two cell lines.

Finally, we assessed NMIIB-S1 TSMod, which was designed to report on the tension generated by a single head of NMIIB. HEK293 cells expressing NMIIB-S1 TSMod yielded an average mTFP1 lifetime of 2.2 ± 0.1 ns, corresponding to a FRET efficiency of 22 ± 4% (mean ± S.D.) (Fig. S4, B, B’ and D). The higher average FRET efficiency relative to NMIIB-S1 TSMod is consistent with a lower degree of average tension. An NMIIB-S1 TSMod dark acceptor mutant served as a control construct and yielded low FRET efficiency, as anticipated (Fig. S4).

To evaluate the possibility that FRET values observed for NMIIB TSMod might be influenced by intermolecular FRET within NMII bipolar filament assemblies, we cotransfected HEK293 cells with NMIIB-TSMod T and the NMIIB-TSMod V, which lack functional FRET acceptor and donor fluorophores, respectively (Fig. S5). FLIM-FRET measurements yielded a FRET efficiency of 2 ± 5% (mean ± S.D.) (Fig. S5), indicating that intermolecular FRET was negligible.

### Inhibition of NMII motor activity significantly increases the FRET efficiency of NMIIB TSMod

We performed FLIM measurements on HEK293 cells expressing NMIIB TSMod before and after treatment with S-nitro-blebbistatin, which inhibits myosin motor activity directly, or ML-7, which downregulates myosin contractility by inhibiting its activating kinase myosin light chain kinase (MLCK) (Kovacs et al., 2004; Lucas-Lopez et al., 2008; Saitoh et al., 1987). The FRET efficiency for NMIIB TSMod increased to 27 ± 1% (mean ± S.D.) in blebbistatin treated cells (Fig. 3B, B’ and I) compared to the 18 ± 3% (mean ± S.D.) recorded before blebbistatin treatment (Fig. 3A, A’ and I). ML-7 treatment led to a comparable increase in FRET efficiency to 28 ± 3% (mean ± S.D.) (Fig. 3C, C’ and I). We also generated a motor domain mutant, NMIIB TSMod R709C, that decreases the maximal ATPase activity 3-fold and abolishes the ability of the mutant myosin to translocate actin (Kim et al., 2005). Cells expressing NMIIB TSMod R709C exhibited an average FRET efficiency of 25 ± 4% (mean ± S.D.), which was significantly increased, compared to cells expressing NMIIB TSMod (p value <0.0001, mann-whitney test) (Fig. 3D, D’, J). Acute inhibition of NMII ATPase activity using S-nitro-blebbistatin or ML-7 did not alter the FRET efficiencies for NMIIB TSMod R709C (Fig. 3D-E, J). Together, these data indicate that FRET values reported by NMIIB TSMod are sensitive to perturbations that alter force generation by NMII.

**Figure 3:**
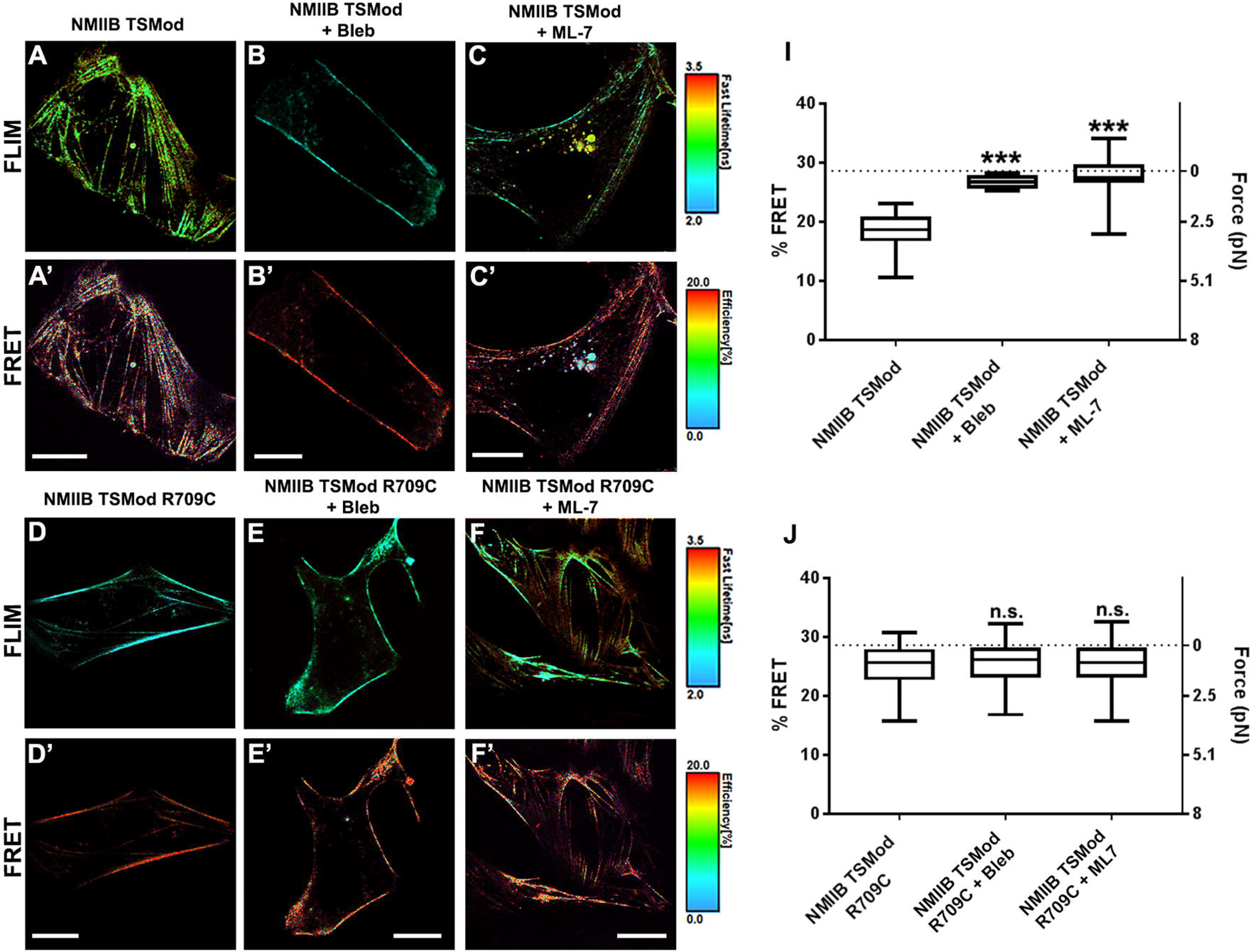
Inhibition of motor activity increases FRET efficiency for NMIIB TSMod. Representative images from time correlated FLIM (A-C) and the corresponding FRET percentages (A’-C’) for HEK293 cells expressing NMIIB TSMod before (A,A’) and after treatment with blebbistatin (B,B’) or ML-7 (C,C’). ROI measurements performed along actomyosin filaments in live cells expressing NMIIB TSMod show decrease in force (pN) along actomyosin filaments as reported by the FRET percentages after 15 min treatment with S-nitro (-) blebbistatin (B’, I) or ML-7 (C’, I). Statistical analysis using Kruskal-Wallis test with multiple comparisons shows significant differences (P value <0.0001). Cells expressing the motor domain point mutant NMIIB TSMod R709C show increased FRET efficiency (D’, J) compared to NMIIB TSMod (A’, I). Treatment with S-nitro (-) blebbistatin (E’,J) or ML-7 (F’, J) did not yield statistically significant increases in FRET efficiency. n = 15-20 cells, ROI FLIM-FRET measurements extracted from point measurements and whole cell images using PicoQuant SymphoTime software were pooled from three separate experimental sets. Dotted line indicates the unloaded FRET efficiency for TSMod (Grashoff et al., 2010), Scale: 10 μm.

### NMIIB TSMod detects mechanical heterogeneity in actomyosin-associated force during cell migration

NMII-mediated force generation helps drive focal adhesion dynamics, retrograde actin flow, and actin treadmilling, all of which are essential for cell migration (Pasapera et al., 2015; Yam et al., 2007). As part of our characterization of NMIIB TSMod, we expressed it in Madin-Darby Canine Kidney (MDCK) cells, a standard model cell line for epithelial biology. FLIM-FRET measurements yielded an average FRET efficiency of 20 ± 5% compared to 10 ± 3% (mean ± S.D.) recorded by mTFP1-NMIIB (Fig. S6). Non-zero FRET measured for the control construct may reflect modestly different intrinsic lifetime for mTFP1 in MDCK cells. Interestingly, stable expression of NMIIB TSMod in MDCK cells created a highly migratory phenotype; these cells migrated faster than the U2OS and HEK293 stable cells expressing NMIIB TSMod.

We took advantage of this observation and used MDCKs as a model system in which to study spatial and temporal variations in tension on NMIIB in migrating cells (Movies 1&2). ROI analysis of actomyosin filaments in the cell periphery, underneath the center of the cell, and the posterior trailing edge of migrating cells showed marked differences in FRET efficiencies (Fig. 4B-D). Converted to implied force values using a published calibration curve (Grashoff et al., 2010), these FRET percentages translate to force magnitudes ranging from ~0.5 − 4 pN (Fig. 4C). We observed marked fluctuations in average FRET efficiencies for all three regions, suggestive of dynamic fluctuations in myosin force generation (Fig. 4D).

**Figure 4:**
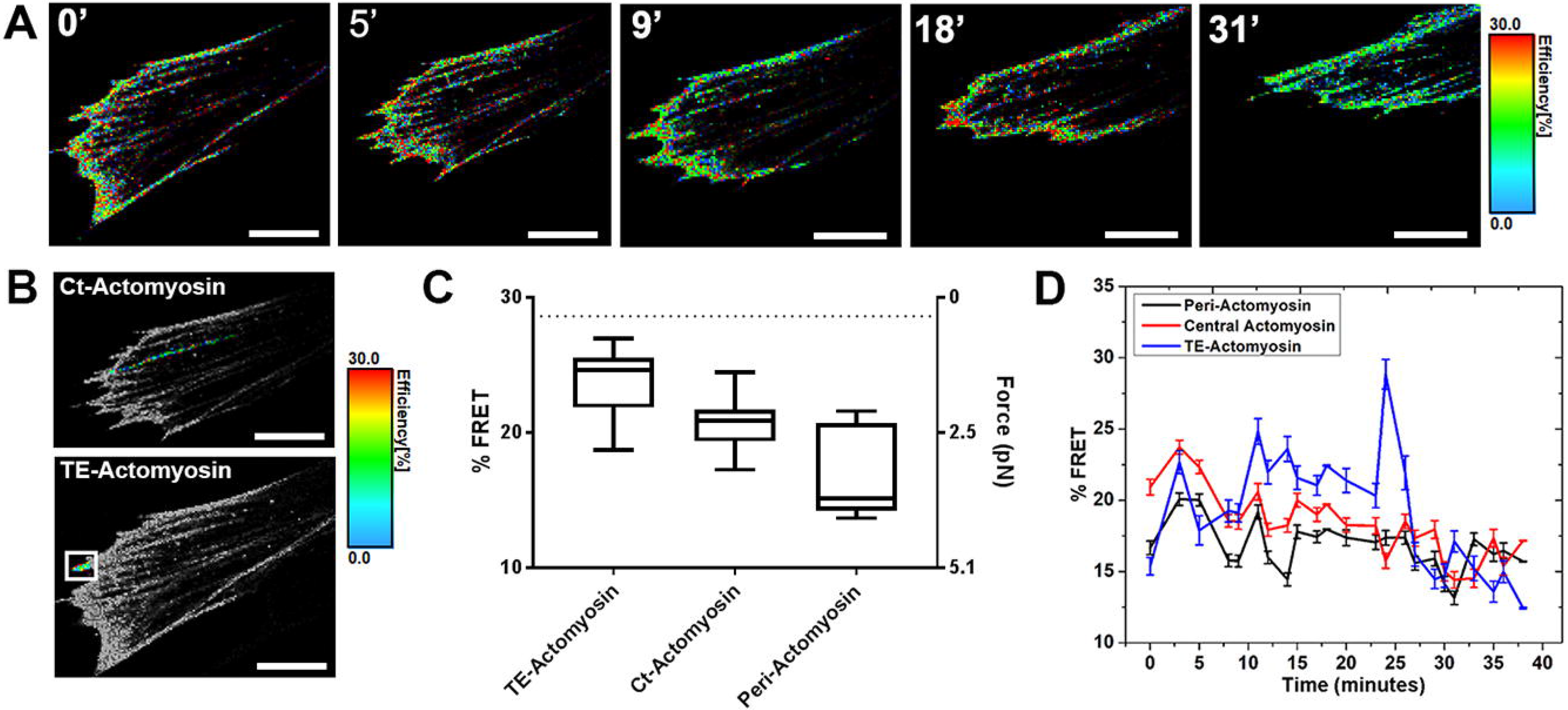
Mechanical heterogeneity detected using NMIIB TSMod during cell migration. Time-course FRET images of a migrating MDCK cell (A). Images show highlighted ROI’s of a central actomyosin fiber, and actomyosin in the trailing edge of a migrating cell (B). Variations in FRET efficiencies detected for multiple ROI’s along peripheral (peri), central (Ct) and trailing edge (TE) actomyosin filaments (C). The time-course of FRET efficiency values detected along peripheral, central actomyosin, and trailing edge associated actomyosin of the migrating MDCK cell show dynamic variations over time, Error bars were calculated by dividing FRET efficiency by the number of photon events (D). Scale: 10μm.

### Discussion

We developed and validated a NMII tension sensor that yielded the first, to our knowledge, direct measurements of forces generated by NMII in living cells. We validated NMIIB TSMod using time-domain FLIM in three different cell types with proper controls to account for the possible contribution of environmental sensitivity, intermolecular FRET, and ATPase activity (mTFP NMIIB; NMIIB TSMod T and V, NMIIB TSMod R706C). In addition, we tested the insertion of TSMod at two locations in NMIIB and found that an insertion point a short distance into the coiled-coil domain was optimal in terms of sensitivity. Measurements in which myosin activity was reduced using either blebbistatin or ML-7 yielded FRET efficiencies in good agreement with the expected FRET values for TSMod under negligible tension (Fig. 3A-C, 3I) (Grashoff et al., 2010; Rothenberg et al., 2018). Interestingly, NMIIB TSMod exhibited subcellular variations in FRET efficiency suggestive of local variation in force generation by NMIIB (Fig. 4). Exploring the origins of these variations lies outside the scope of this report. We note though that our measurements are consistent with FRET values reported for a vinculin tension sensor in the front (low FRET) and rear (high FRET) of migrating endothelial cells (Grashoff et al., 2010), a correspondence that provides additional evidence that NMIIB TSMod reports on localized cytoskeletal tension.

The R709C mutation is known to decrease NMIIB actin-activated ATPase activity to 30% that of wild-type NMIIB, and to completely block actin translocation in an *in vitro* motility assay (Kim et al., 2005). Consistent with these observations, the crystal structure of NMIIB indicates that this mutation likely blocks allosteric communication between the ATPase site and the lever arm (Munnich et al., 2014). Interestingly, the R709C motor domain mutant was sufficient to rescue cytokinesis defects in both NMIIB-depleted COS-7 cells and cardiac myocytes (Ma et al., 2012); an observation interpreted as indicating that NMIIB R709C could maintain tension, but not generate active force. In agreement with this, we observed a FRET efficiency for NMIIB R709C of 25 ± 3.6, suggestive of the presence of a small amount of tension on NMIIB TSMod R709C molecules. Consistent with previous interpretations, this tension (if present) is unlikely to require the ATPase activity of the mutant myosin, as treatment with S-nitro-blebbistatin or ML-7 did not alter the FRET efficiencies for NMIIB TSMod R709C.

It is likely that further optimization will yield myosin-based tension sensors with expanded sensitivity and applicability. New sensor modules with improved molecular springs may improve the sensitivity of the current sensor designs (LaCroix et al., 2018; Ringer et al., 2017). The current sensor design can be likewise extended to NMIIA, the other major NMII isoform in most cells. Complementary NMIIA and NMIIB sensors will likely be useful in clarifying how the two isoforms differentially regulate cytoskeletal organization and force generation in a variety of physiological contexts (Murrell et al., 2015; Vicente-Manzanares et al., 2011). Finally, it will be of considerable interest to translate the current sensor designs to model organisms, for example *Drosophila* and *C. elegans,* to directly visualize myosin-driven forces during embryogenesis.

## Materials and Methods

### Generation of NMIIB tension sensors and control constructs

Infusion Cloning (Clontech cat.# 638910) was used for DNA manipulation unless otherwise specified. Human MYH10 (NMIIB) coding sequence was amplified using an NMIIB-GFP plasmid generated in the laboratory of Dr. Robert Adelstein, as used in our previous work (Chandrasekar et al., 2014). The NMIIB coding sequence was amplified with primers *PCDNA3IIB FL F* and *PCDNA3IIB FL R* and inserted between EcoR1 and Apa1 of the PCDNA3 multiple cloning site to generate NMIIB-pCDNA3. A unique XBA1 restriction site beginning at base pair 3663 in the NMIIB coding sequence was used to linearize the NMIIB-pCDNA3 construct. The TSMod Coding sequence was amplified from Addgene Plamsid # 26021 (Schwartz Lab). Primers *TSMod XBA1 F* and *TSMod XBA1 R* were used to amplify the TSMod sequence and it was inserted into the Xba1 linearized NMIIB-pCDNA3 construct to generate the NMIIB TSMod construct.

NMIIB TSMod Control Constructs. mTFP1-NMIIB was generated by amplification of NMIIB coding sequence using primers *ICCP170* and ICCP171 inserted into mTFP1-NMIIA (Addgene #55501) replacing NMIIA. Constructs were verified by sequencing analysis.

Three NMIIB TSMod mutant constructs were generated for validation purposes. NMIIB TSMod T (G73S), harboring a point mutation that renders mTFP1 inactive, NMIIB TSMod V (G67S) which renders the Venus fluorophore inactive, and NMIIB-R709C (NMIIB motor domain mutant). Point mutants were generated using the NEB Q5 Site directed mutagenesis kit and protocol (New England Biolabs #E0552S). NMIIB TSMod T (G73S) was generated using primer pair *ICCP269* and *ICCP270*. NMIIB TSMod V (G67S) was generated using primer pair *ICCP304* and *ICCP305*. NMIIB-R709C TSMod was generated using primer pair *ICCP259\** and *ICCP260\**. To generate NMIIB-5AA, the 40 residue molecular spring in the TSMod was replaced with a 5 amino acid SGKRS linker between the mTFP1 and Venus fluorophores. The NEB Q5 kit polymerase and protocol was used with primers *ICCP294* and *ICCP295* to inversely amplify the entire NMIIB-TSMod plasmid sequence lacking the 40 residue molecular spring region. Homologous 15bp overhangs on the 3’ end of the inverse PCR primers were ligated following the NEB T4 Ligation protocol (New England Biolabs # M0202S) to introduce the 5AA sequence. Constructs were verified by sequencing analysis. (*Note all primers ordered from Eurofins Genomics referenced in methods are located in Table 1*).

**Table 1:**
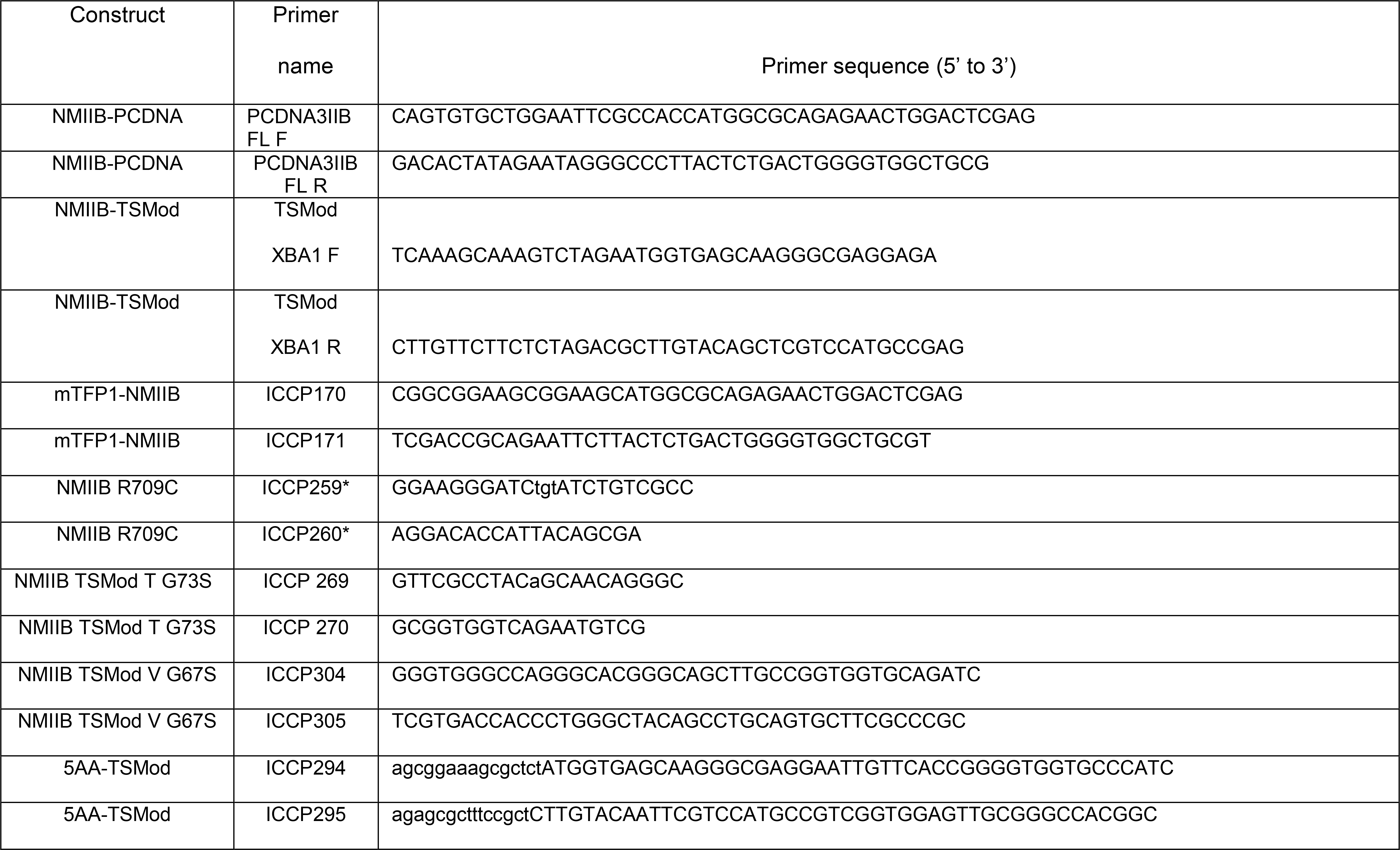

NMIIB-S1 TSMod was generated as described below. The human coding sequence for MYH10 gene was derived from Addgene plasmid ID# 11348 (Wei & Adelstein, 2000). The NMIIB coding sequence was then inserted into pEGFP-N3 between the KpnI and NotI sites, which also removed the parent vector’s GFP tag. TSMod (Grashoff *et al.,* 2010; Borghi *et al.,* 2012) was cloned into the NMIIB sequence at the beginning of the rod tail domain (between L845 and Q846) using BspEI and SpeI cloning sites engineered into the sequence. We used a TSMod variant that uses mTFP1 and mEYFP as the FRET donor and acceptor respectively, attached to the NMIIB coding sequence by short linkers encoding GlyGlyAlaGlyAlaGly to aid in fluorophore folding. Additionally, PacI and AscI restriction sites between the fluorophores and the flagelliform sequence were inserted to aid in the construction of control constructs. The NMIIB-S1 TSMod domain sequence is thus NMIIB (catalytic domain and lever arm)-linker-BspEI-mTFP1-PacI-flagelliform-AscI-mEYFP-SpeI-linker-NMIIB (coiled coil domain). A dark acceptor version of NMIIB-S1 TSMod was made by introducing the mutation Y67G (relative to the fluorophore sequence) in mEYFP (Takanaga & Frommer, 2010).

### Cell Culture Conditions and Stable Cell Generation

HEK293, U2OS, Mouse Embryonic Fibroblast (MEF) WT and MEF NMIIB KO cells were cultured in DMEM (Corning #10013CV) supplemented with 10% FBS (Atlanta Biologicals# S11150H) and Penicillin/Streptomycin (Corning #30002Cl). MDCK cells were cultured in EMEM (Corning #10009CVR) supplemented with 10% FBS. Cells were cultured at 37 °C with 5% CO_2_. Lipofectamine 2000CD (Life Technologies #12566014) with the corresponding protocol was used to transfect the cell lines. To generate cells stably expressing these constructs, media supplemented with 500 μg/mL G418 Sulfate Solution (Corning #30234CR) was used on cells two days after transfection. After a ten-day G418 selection, GFP positive cells were sorted using the FACSjazz cell sorting system (BD Biosciences) to obtain a pure population. G418 selection pressure was maintained while culturing cells.

### Immunocytochemistry

35mm Fluorodish glass bottom dishes (World Precision Instruments #FD35-100) were coated overnight with human fibronectin (Corning #CB 40008). Each dish was rinsed with sterile phosphate-buffered saline (Corning# 21040CV) prior to seeding cells on dishes. 150,000 cells were seeded onto each dish for immunocytochemistry. 18 to 24 hours after seeding, cells were transfected as described above and prepared for immunocytochemistry. Cells were fixed for 15 minutes at room temperature with 4% paraformaldehyde (Electron Microscopy Sciences #15710) in cacodylate buffer (0.1 M Cacodylate Salt, 10 mM CaCl_2_, 10 mM MgCl_2_, pH 7.4) The solution was warmed to 37 °C prior to addition to cells. Cells were rinsed 3x with cacodylate buffer followed by PBS. Cells were permeabilized by incubating in 0.1% Triton-X-100 in PBS for 10 minutes. Blocking solution (5% goat serum, 4% fish gelatin, 1.2% BSA in phosphate buffered saline) was applied for 1 hour at room temperature. After blocking, primary Anti-NMIIB Rabbit antibody (Biolegend #909901) was applied at a 1:500 dilution for 1 hour at room temperature. Secondary antibodies and dyes used were Anti-Rabbit Alexa Fluor 567 (Life Technologies #A-11011) at a 1:500 concentration and Alexa Fluor 647 Phalloidin at a 1:100 concentration (Life Technologies #A22287). Each secondary antibody was incubated with cells for 1 hour at room temperature. After final PBS washes, cells were mounted using Vectashield with DAPI (Vector Labs #H-1200).

### Western Blot Analysis

HEK293 cells transfected with NMIIB TSMod and mTFP1-NMIIB were lysed 48 hours after transfection. Cells were lysed with ice cold NMIIB Lysis Buffer (150 mM NaCl, 50 mM Tris Base, 5 mM MgCl_2_, 2 mM ATP, 0.1% NP-40, 10% Glycerol). After lysis, protein levels were quantified using the colorimetric BCA assay (Pierce #23225). 10 μg of protein lysate was loaded on a 5% tris glycine polyacrylamide gel and protein separation occurred at 110 volts in 25 mM Tris, 192 mM glycine, and 0.1% SDS running buffer (pH 8.3). Proteins were transferred to PVDF membranes at 100 volts for 1 hour in ice cold 25 mM Tris, 192 mM Glycine transfer buffer (pH 8.3). Membranes were rinsed briefly in Tris-buffered saline with Tween 20, TBST (50 mM Tris HCl, 150 mM NaCl, pH 7.5 0.1% Tween-20). Membranes were blocked in 5% milk TBST for one hour at room temperature with agitation. Anti-NMIIB Rabbit (Biolegend #909901) primary antibody was diluted 1:1000 in 5% milk TBST and placed on membrane overnight in a rocker platform at 4 °C. Membranes were rinsed three times with TBST. Anti-Rabbit HRP secondary antibody (Jackson Immunoresearch #711-035-152) was diluted 1:10,000 in 5% milk TBST and incubated with the membrane for one hour at room temperature with gentle agitation. The membrane was then rinsed three times with TBST and visualized with UV lighting after applying the chemiluminescent substrate to the membrane (EMD Millipore #WBLUF0100).

### Confocal Imaging

Imaging was performed with a Nikon A1-TIRF microscope (Sanford Research) and the Olympus Fluoview 1000 microscope (IUPUI).

### FRET imaging

Fluorescence lifetime images were acquired by a custom-made fluorescence lifetime imaging microscope built on a laser scanning confocal microscope (FluoView 1000, Olympus). A picosecond pulsed laser with an excitation wavelength of 450 nm (LDH-D-C-450, Picoquant) was coupled with the laser scanning module, and the fluorescent signal was filtered by a band-pass filter centered at 490 nm (ET490/40X, Chroma) before entering a photon counting detector (PD-100-CTC, MPD). All signals were recorded in Time-Correlated Single Photon Counting (TCSPC) mode with a data acquisition board (TimeHarp 260, Picoquant). The FRET efficiency μ was calculated based on the lifetime of the donor molecule:

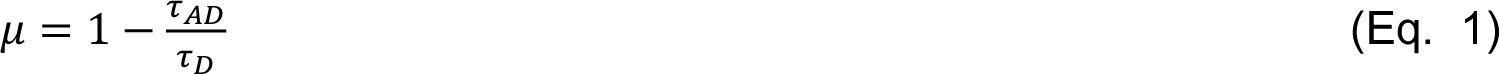

where *τ*_*AD*_ is the fluorescence lifetime of the donor molecule in the presence of the acceptor, and *τ*_*D*_ is the fluorescence lifetime of the donor molecule without the acceptor. The value of *τ*_*AD*_/*τ*_*D*_ was obtained by fitting the decay curve in the software Symphotime (Picoquant) (Jing Liu, 2014; Vidi et al., 2014). We estimated the tensile force using FRET efficiency as per Grashoff *et al* (Grashoff et al., 2010).

### Statistical analysis

3-5 independent experimental replications were conducted for each experiment (Fig. 2 – 4). Statistical significance was evaluated using one-way analysis of variance (ANOVA), as described in figure legends.

## Supporting information

MDCK Movie 1

MDCK Movie 2

---

**Figure S1:**
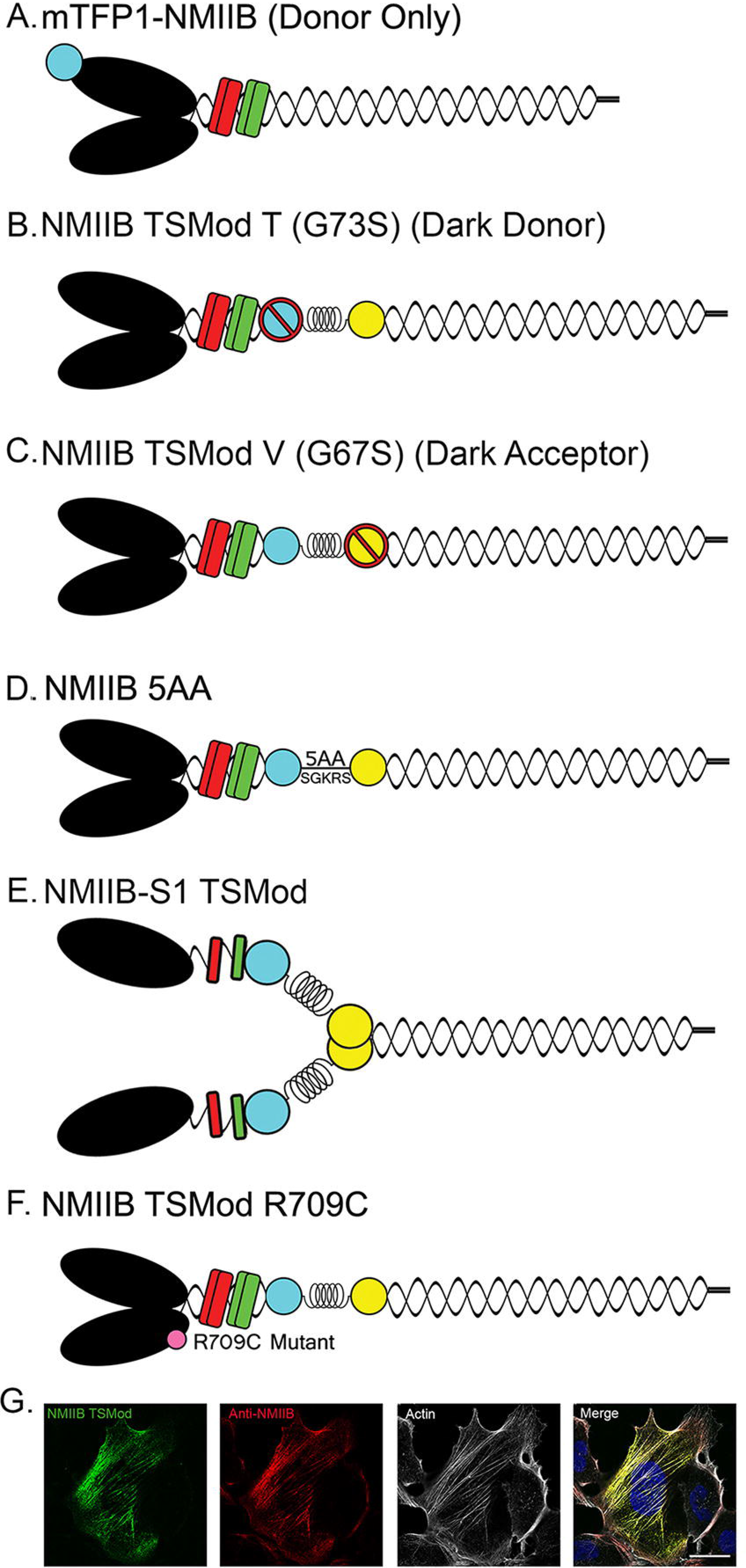
Cartoons illustrating the various NMII constructs, Expression of NMIIB tension sensor. Cartoon illustrating the structural domains of various NMIIB constructs (A-F). HEK293 cells transfected with NMIIB TSMod show NMIIB TSMod (green, 488nm) localizing with endogenous NMIIB (red, cy3) and F-actin (white, cy5) in fixed and immunostained cells (G).

**Figure S2:**
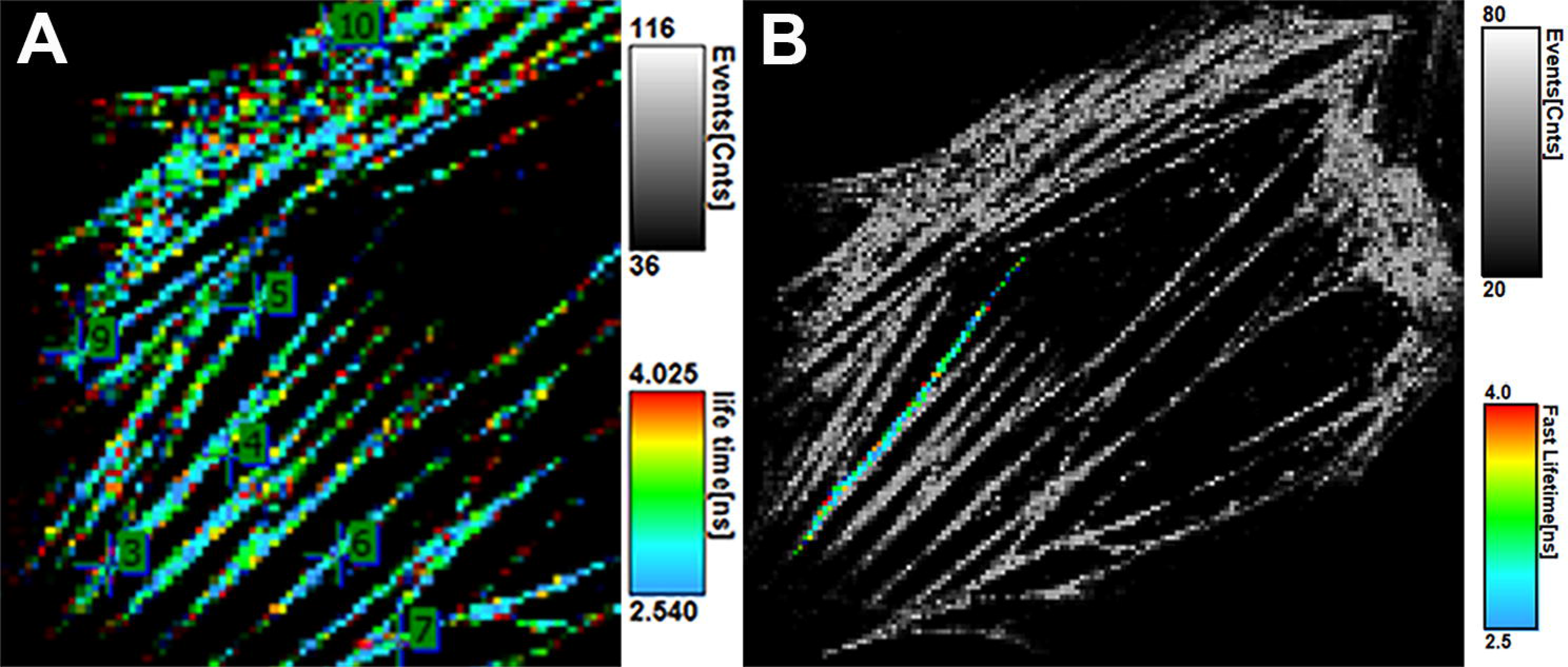
ROI FLIM-FRET measurement strategy. Representative images from time correlated FLIM (A,B) of a HEK 293 cell expressing NMIIB TSMod. ROI based point FLIM measurements (points 3-7, 9, 10) were performed on randomly selected regions from multiple actomyosin filaments in a living cell (A). 10 – 30 ROIs were selected per cell depending on the size of the cell and the number of stress fibers. (B) Example ROI with FLIM values extracted for a highlighted actomyosin filament using PicoQuant Symphotime software (B). Number of events and lifetime (ns) are indicated alongside the images.

**Figure S3:**
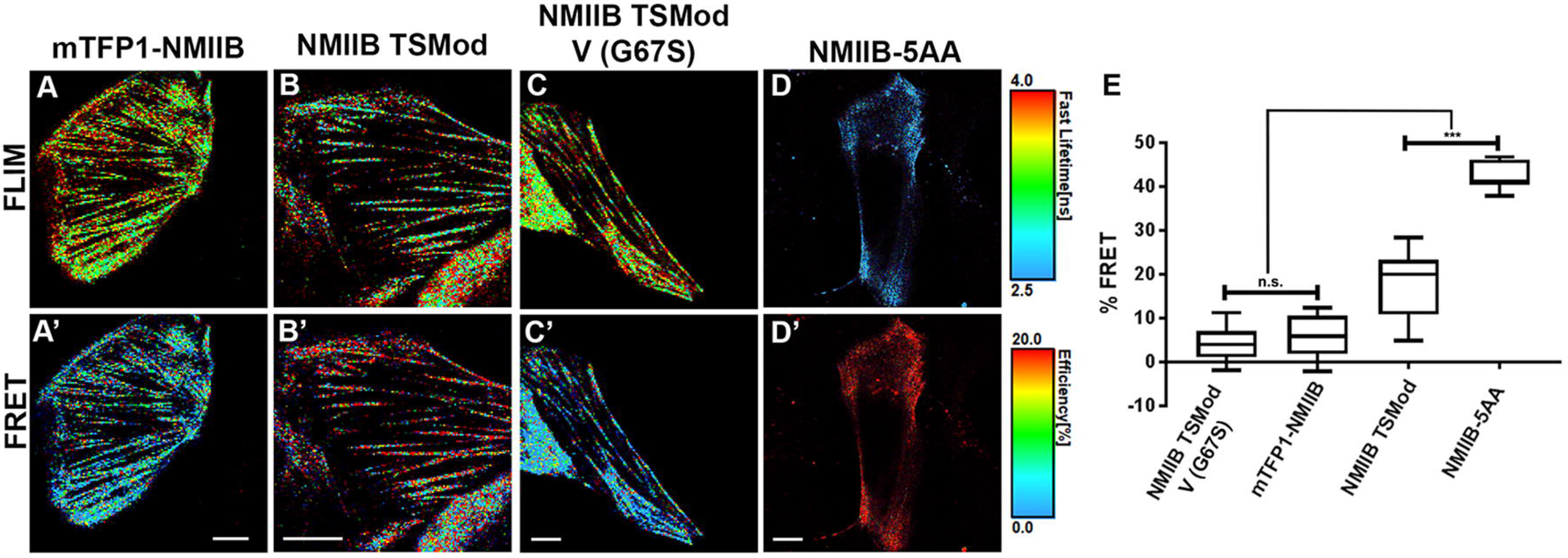
NMIIB TSMod FLIM-FRET measurements in U2OS osteosarcoma cells. Representative images from FLIM (A-D) and the corresponding FRET percentages (A’-D’) of U2OS cells expressing the NMIIB TSMod and the control constructs. Scale: FLIM (2.5 – 4 ns) and FRET (0 – 20%). ROI measurements performed along actomyosin filaments in live cells expressing mTFP1-NMIIB [donor only control (A, A’, and E)] and the NMIIB TSMod V (G67S) [dark acceptor control (C, C’ and E)] show lower FRET efficiencies (%) compared to NMIIB TSMod (B, B’, E). NMIIB-5AA, a FRET control that measures energy transfer between fluorophores at close proximity, shows low lifetimes and high FRET efficiencies (D, D’ and E). n = 15-20 cells. ROI FLIM-FRET measurements extracted from point measurements and whole cell images using PicoQuant SymphoTime software were pooled from 3-5 separate experimental sets. (P value < 0.0001) between all parameters, except between controls NMIIB TSMod G67S and mTFP1-NMIIB, Scale: 10 μm.

**Figure S4:**
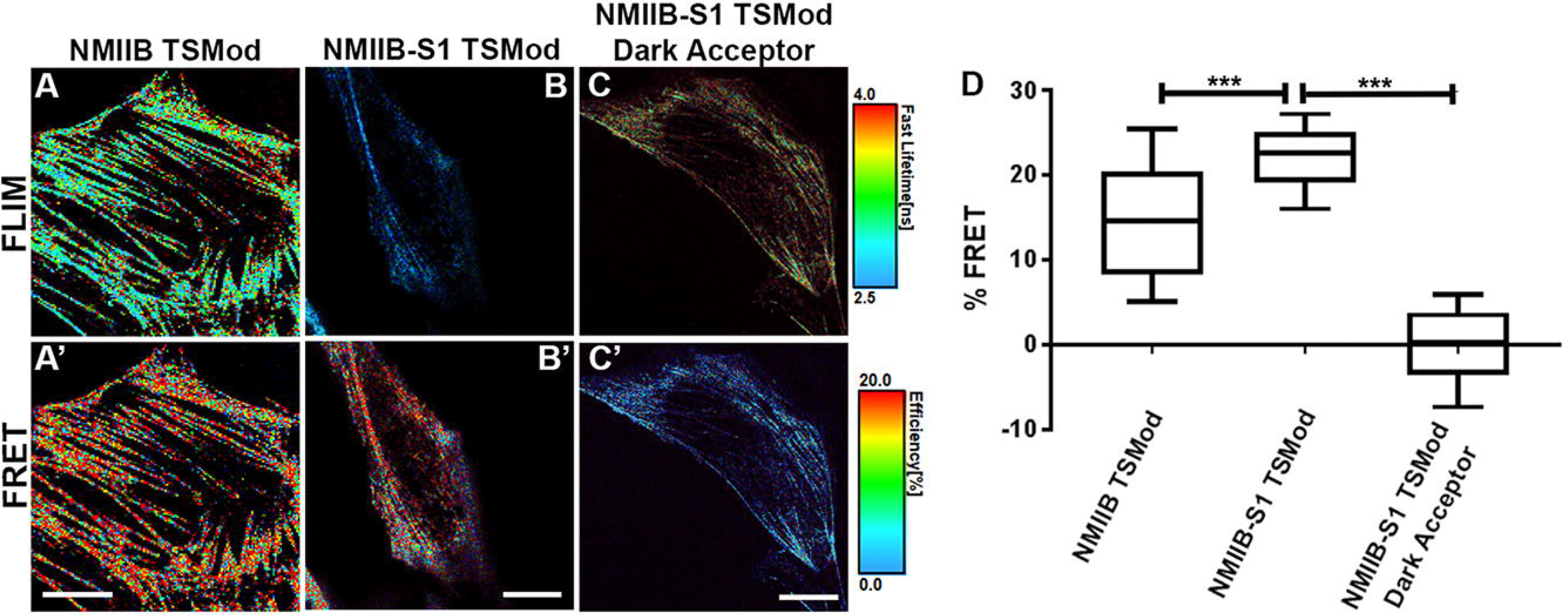
NMIIB-S1 TSMod reports a narrow range of FRET values compared to NMIIB TSMod. Representative images from time correlated FLIM (A-C) and the corresponding FRET percentages (A’-C’) for HEK293 cells expressing the NMIIB TSMod, NMIIB-S1 TSMod and NMIIB-S1 TSMod dark acceptor control construct, Scale: FLIM (2.5 – 4 ns) and FRET (0 – 20%). ROI measurements performed along actomyosin filaments in live cells expressing NMIIB TSMod (A, A’, and D) show lower average FRET values, consistent with higher tension, compared to NMIIB-S1 TSMod (B, B’, D). The dark acceptor control construct recorded low FRET efficiency percentages (C, C’, D). n= 15-20 cells for NMIIB TSMod and 10 cells for NMIIB-S1 TSMod and NMIIB-S1 TSMod dark acceptor. FRET measurements were pooled from 3-5 separate experimental sets. Statistical analysis using Kruskal-Wallis test with multiple comparisons produced a P value of <0.0001 between NMIIB TSMod and NMIIB-S1 TSMod, Scale: 10 μm.

**Figure S5:**
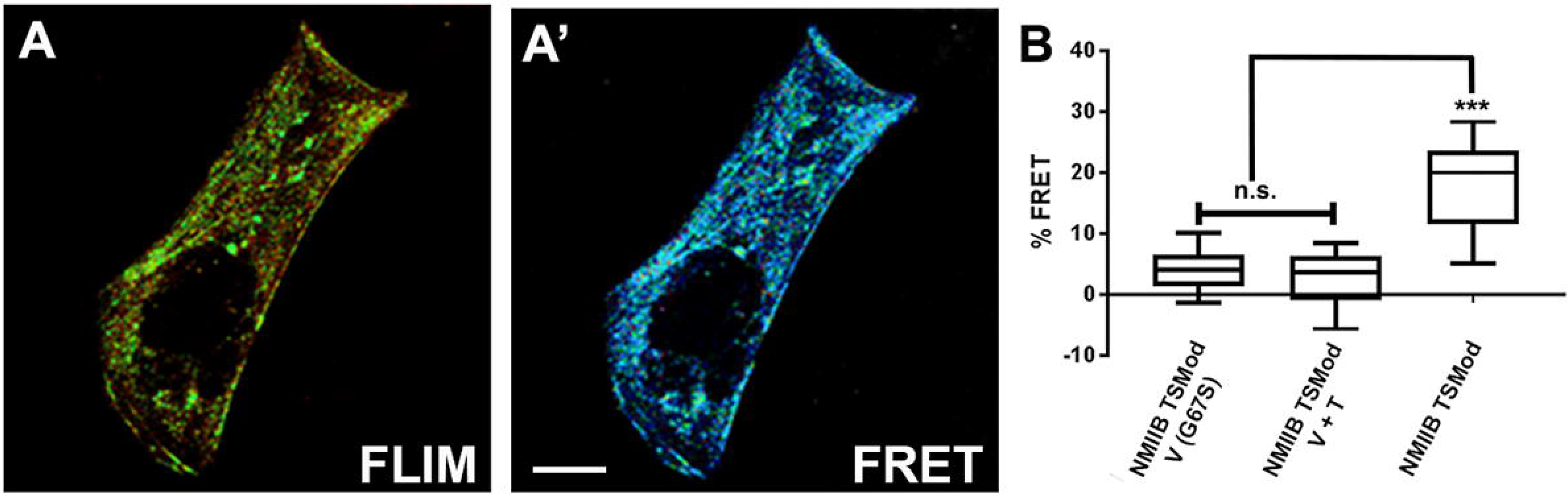
Intermolecular FRET is negligible for NMIIB TSMod. Representative FLIM (A) and FRET (A’) images show HEK 293 cells co-expressing NMIIB TSMod V (G67S) (dark acceptor) and NMIIB TSMod T (G73S) (dark donor). Point measurements performed along the actomyosin filaments in live cells co-expressing both constructs show low FRET efficiency percentages compared to NMIIB TSMod (B) indicating negligible intermolecular FRET. Statistical analysis using Kruskal-Wallis test with multiple comparisons shows significant differences (P value < 0.0001) between NMIIB TSMod and NMIIB TSMod G67S as well as co-expressing NMIIB TSMod G67S V + T, but not between the controls (P value = 0.7882). Scale: 10 μm.

**Figure S6:**
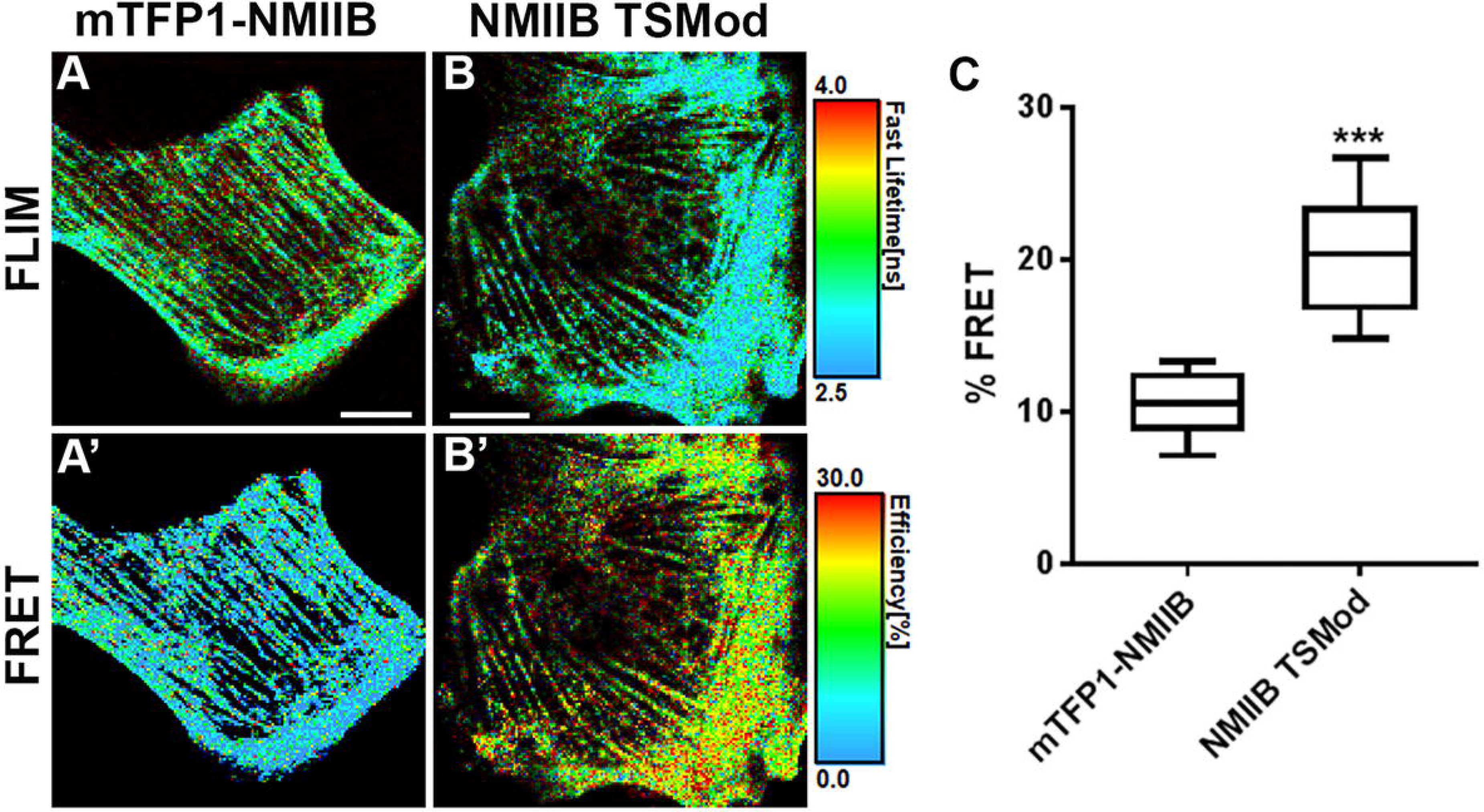
NMIIB TSMod FLIM-FRET measurements in MDCK cells. Representative images from time correlated FLIM (A,B) and the corresponding FRET percentages (A’, B’) of MDCK cells expressing the NMIIB TSMod and the donor only control mTFP1-NMIIB. Scale: FLIM (2.5 − 4 ns) and FRET (0 − 30%). ROI measurements performed along actomyosin filaments in live cells expressing mTFP1-NMIIB [donor only control (A, A’, and C)] show lower FRET % compared to NMIIB TSMod (B, B’, C). ROI FLIM-FRET measurements extracted from point measurements and whole cell images using PicoQuant SymphoTime software were pooled from three separate experimental sets. (P value <0.0001) between both parameters using Mann-Whitney test. Scale: 10 μm.

